# Striatal indirect pathway mediates hesitation

**DOI:** 10.1101/2024.09.16.613332

**Authors:** Matthew A Geramita, Susanne E Ahmari, Eric A Yttri

## Abstract

Determining the best possible action in an uncertain situation is often challenging, and organisms frequently need extra time to deliberate. This pause in behavior in response to uncertainty – also known as hesitation – commonly occurs in many aspects of daily life, yet its neural circuits are poorly understood. Here we present the first experimental paradigm that reliably evokes hesitation in mice. Using cell-type specific electrophysiology and optogenetics, we show that indirect, but not direct, pathway spiny projection neurons specifically in the dorsomedial striatum mediate hesitation. These data indicate that the basal ganglia circuits controlling the pausing involved in cognitive processes like hesitation are distinct from those that control other types of behavioral inhibition, such as cue-induced stopping.

## Introduction

Hesitation – the pausing of action in the face of uncertainty^1^ – is an important cognitive process used in a variety of real-world scenarios. It is a critical component of deliberation^2,3^ and provides the crucial additional time needed to formulate a thoughtful answer to a delicate question or to decide whether to answer a phone call from an unknown number. While hesitation is often beneficial, deficits in hesitation can become pathological and contribute to psychiatric symptoms such as impulsivity and anxiety. Despite the ubiquity of hesitation in daily life and its importance in decision-making^2,3^, little is known about its underlying neural circuitry in large part because current behavioral paradigms fail to isolate the key aspects of hesitation that make it distinct from other scenarios in which one must inhibit an action. For example, stop-signal tasks (SSTs) have been invaluable in developing ‘race models’ to conceptualize behavioral inhibition^4–8^. In these models, separate Stop and Go processes, represented respectively in the activity of the hyperdirect pathway input to the subthalamic nucleus^5,9^ and the striatal direct pathway, race to determine whether an action is carried out or successfully aborted after the presentation of a stop-signall^10^. However, hesitation is a fundamentally different behavior than the stopping measured during SSTs. The decision to hesitate is internally generated, rather than a response to an explicit cue, and requires significant cognitive control to identify uncertain contexts in which pausing is beneficial. Therefore, an open question is whether hesitation is regulated by the same neural circuits that mediate cue-induced stopping.

To resolve this question, we developed a novel experimental procedure that reliably evokes hesitation in mice. Using an auditory association task, we show that mice hesitate – that is they pause anticipatory licking exclusively in response to cues associated with an uncertain reward. We then discovered that hesitation is mediated by a specific population of dorsomedial striatum (DMS) indirect pathway neurons. We performed cell-type specific *in vivo* electrophysiology and found a population of indirect pathway spiny projection neurons (iSPNs) in the DMS that signal uncertain tone-reward associations. Additionally, the degree of hesitation correlated with trial-to-trial changes in the activity of these uncertainty-responsive iSPNs. Finally, we showed that increasing iSPN activity increased hesitation, while decreasing iSPN activity decreased hesitation. Importantly, neither DMS dSPNs, nor dSPNs or iSPNs in a separate striatal region influenced hesitation, suggesting a specific role for DMS iSPNs in hesitation. Taken together, we show that the indirect pathway of the basal ganglia plays a critical role in hesitation, a poorly understood but important behavior in both health and disease, and that separate neural circuits underly action cancellation and the pausing involved in cognitive tasks like hesitation.

## Results

### Mice hesitate during an auditory association task

Mice (n=48 mice; 22 females; 141 sessions, 42,125 trials) were trained on an auditory conditioning task in which each of three tones was associated with a different reward probability: Unrewarded (0%), Rewarded (100%), or Uncertain (50%) (**Fig. 1A**). Tones were presented pseudo-randomly, and tone-reward associations were counterbalanced across animals. We observed that anticipatory licking prior to reward delivery was correlated with reward probability (**Fig. 1B,C**; Unrewarded: 1.2 ± 0.1 Hz; Uncertain: 4.4 ± 0.2 Hz; Rewarded: 5.1 ± 0.1 Hz; p<0.001; One-way ANOVA) indicating that mice learned the value of the cues. Additionally, the latency to begin licking differed by trial type. For both Rewarded and Unrewarded trials, mice started licking within 400 ms of tone onset (**Fig. 1D-F**; first lick latency: Unrewarded = 254 ± 45 ms; Rewarded = 343 ± 84 ms). However, on Uncertain trials, mice paused for hundreds of milliseconds after tone onset (first lick latency: 751 ± 71 ms), indicating that cues associated with an uncertain reward reliably induced a self-initiated pause in anticipatory licking (i.e. hesitation). Importantly, pauses in behavior observed during Uncertain trials were not due to differences in cue value because first lick latencies were short for both Rewarded and Unrewarded trials. Next, in a subset of mice, we determined whether tone predictability influenced hesitation. After first presenting tones pseudorandomly (unpredictable), we modified the task so that tones were presented in blocks of 10 and were therefore predictable. In both the unpredictable and predictable versions of the task, anticipatory licking correlated with tone value (**Fig. S1A,B**). However, mice only demonstrated hesitatation when uncertain tones were unpredictable. Hesitation disappeared when Uncertain trials were predictable (**Fig. S1C,D**). This suggests that hesitation only occurs when an animal unexpectedly encounters a cue that leads to an uncertain reward.

**Figure 1:**
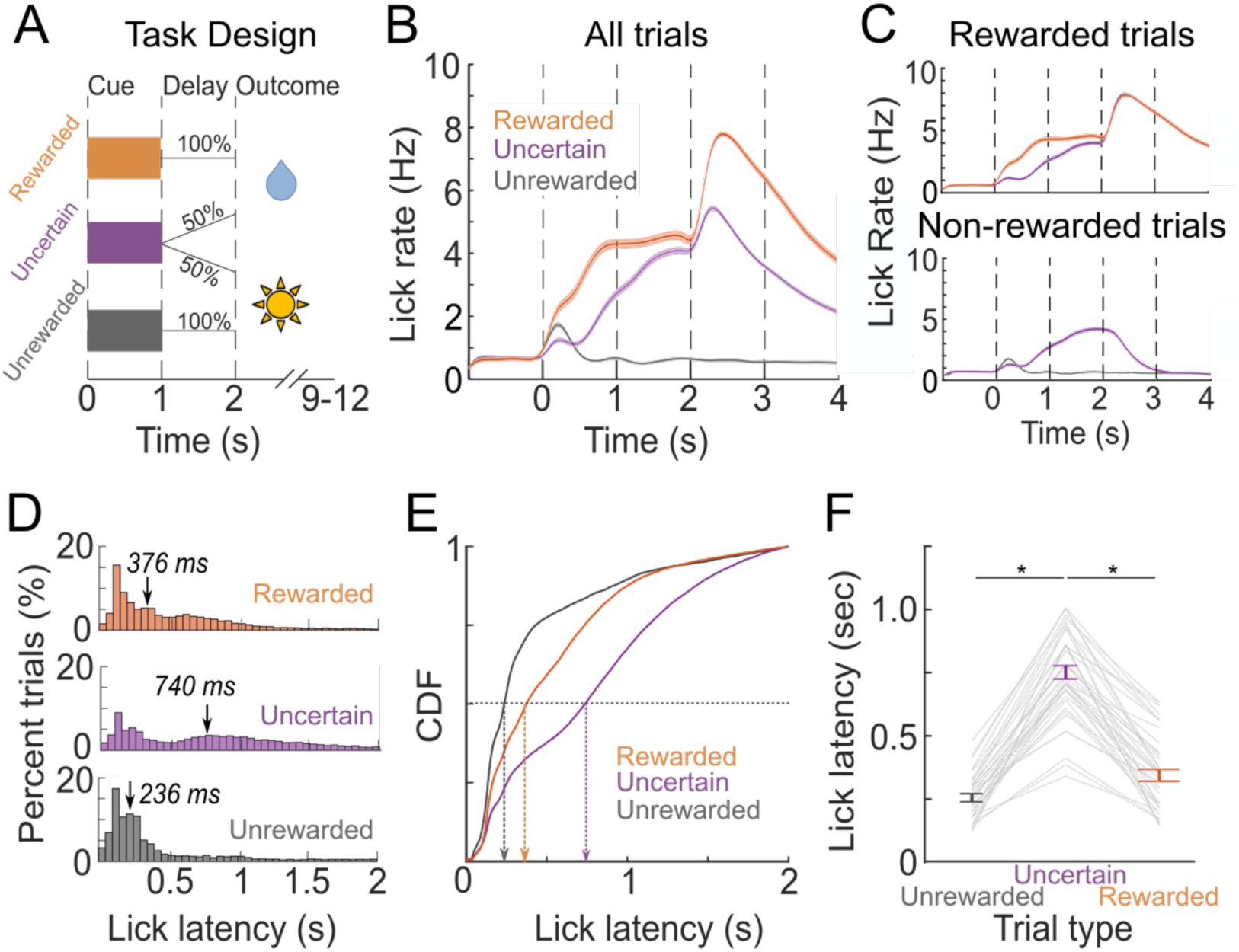
Mice hesitate during an auditory association task. **A**) Task structure: On each trial, one of three tones was played for 1 sec. Following a 1 sec delay, either 4uL drop of sweetened water was dispensed or the house light turned on for 200 ms. Tones were either Rewarded (orange), Unrewarded (grey), or Uncertain (purple). **B-C**) Mice learned the value of each tone, evidenced as the rate of anticipatory licking during the cue and delay periods correlating with the value of the tone (p < 0.001; ANOVA). **B**) Anticipatory licking across all trials. **C**) *TOP:* Anticipatory licking for rewarded trials. *BOTTOM:* Anticipatory licking for unrewarded trials. **D-E**) Mice began anticipatory licking later for uncertain trials compared with unrewarded or rewarded trials. **D**) Probability density function (PDF) of first lick latency for the three trial types. Arrows correspond to the median first lick latency. **E**) Cumulative density function (CDF) of first lick latency. Arrows correspond to the median first lick latency. **F)** Average first lick latency between trials across mice. First lick latency is longer on uncertain trials compared to rewarded or unrewarded trials (p < 0.001; ANOVA). Data taken from 48 mice and 141 sessions for a total of 42,125 trials.

### A population of DMS SPNs preferentially responds to uncertain tones and has activity that covaries with hesitation

How might hesitation be represented in the brain? We hypothesized that for a neuron to mediate hesitation, its activity should have two specific characteristics. First, ‘hesitation neurons’ should preferentially respond to the uncertain tone. Second, trial-to-trial fluctuations in anticipatory licking should co-vary with the activity of these uncertainty-responsive neurons. To test these hypotheses, we used Neuropixels probes^11^ (**Fig. 2A**; see Methods; 9 animals, 24 sessions) to record neural activity from 1047 SPNs (**Fig. S2A**) in the dorsomedial striatum (DMS), an area known to be important for decision-making under uncertainty^12,13^. Next, we isolated 613 SPNs whose activity differed between the three tones and classified them into one of four response types (**Fig. 2B**; see methods): positive value (+Val), negative value (-Val), uncertainty (Unc), or certainty (Cer). Similar to prior reports^14^, a majority (64%) of neurons were modulated by tone value (**Fig. 2C,D**; +Val: 37%, -Val: 27%). However, many SPNs were modulated by tone uncertainty (**Fig. 2E**; Unc: 23%) or certainty (**Fig. 2F**; Cer: 13%), corroborating prior reports that neurons in the striatum signal the uncertainty of tone-reward associations^15^.

**Figure 2:**
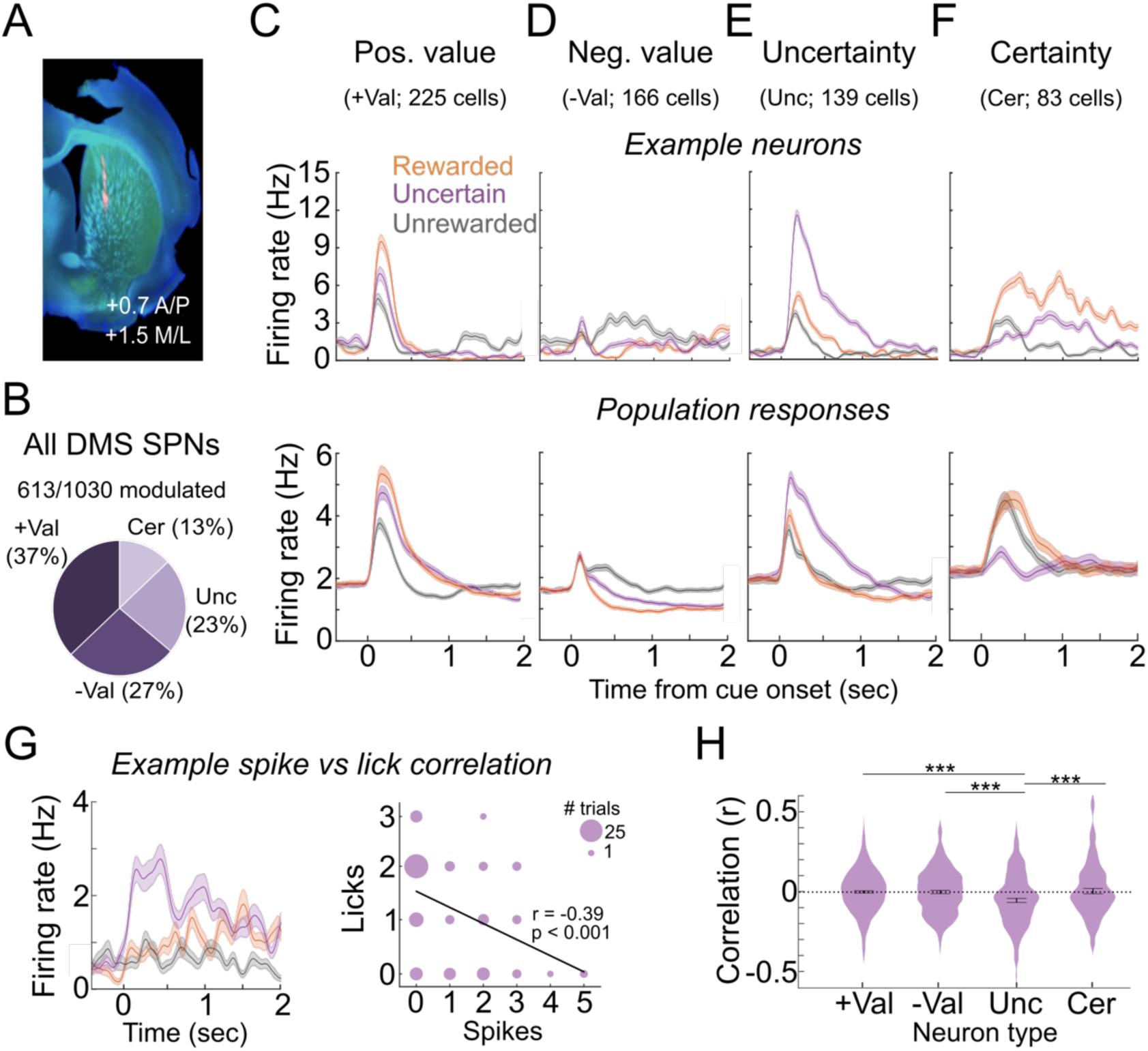
A population of DMS SPNs responds to uncertain tone-reward associations and has activity which covaries with hesitation. **A**) Neuropixels probes recorded neural activity in the DMS and were coated with DiI to confirm recording location. **B**) We recorded from 1030 DMS SPNs (9 mice, 24 sessions) and classified the 613 neurons that were differentially modulated by the tones (see Methods) into one of four groups: positive value (+Val), negative value (-Val), uncertainty (Unc), and certainty (Cer). **C-F**) Example neuron peri-stimulus time histograms (PSTHs) (*Top)* and population average PSTHs (*Bottom)* in each of the four groups: +Val (**C**), -Val (**D**), Unc (**E**), Cer (**F**). **G-H**) To determine whether hesitation correlated with DMS SPN activity, we calculated the spike/lick correlation for each neuron. **G**) *Left:* Example PSTH from an uncertainty-responsive neuron. *Right:* Correlation between spikes and licks for the example neuron calculated within the first 740ms after tone onset (see methods). Marker size indicates the number of trials for each spike/lick combination. **H**) Violin plots of the distribution of spike/lick correlations across the four types of neural responses. The average correlation in the uncertainty-responsive neurons was lower than the average correlations from the other three groups (ANOVA followed by post-hoc t-tests; *** p< 0.01).

We next tested whether the extent of hesitation covaried with SPN activity. To do this, we calculated the correlation between the number of spikes and licks within the first 740 ms of tone onset (i.e. the median first lick latency of uncertain trials) for each trial type and each neuron (**Fig. 2G, Fig. S2B-G**; see methods). On Uncertain (**Fig. 2H**), but not Rewarded or Unrewarded (**Fig. S2B,C**), trials, spike/lick correlations were significantly lower in the uncertainty (r = -0.076 ± 0.0043) population compared to positive value (r = -0.0008 ± 0.01), negative value (r = -0.002 ± 0.02), or certainty (r = 0.004 ± 0.01) populations (p < 0.01, 1-way ANOVA followed by post-hoc t-tests). Consequently, the stronger the activity in the uncertainty-responsive neurons, the fewer licks the animal emits, suggesting that uncertainty-responsive neurons exhibit the neural correlates of hesitation.

### Indirect pathway SPNs are enriched in uncertainty-responsive neurons that correlate with hesitation

A subset of neurons were recorded from mice in which channelrhodopsin (ChR2) was expressed in either A2A+ or DRD1+ neurons, which allowed us to optogenetically identify 99 indirect (iSPNs) and 112 direct (dSPNs) pathway neurons (**Fig. 3A,B**; see methods), respectively. Similar to the overall dataset, a majority of iSPNs (**Fig. 3C**) and dSPNs (**Fig. 3F**); were modulated by tone value. However, more iSPNs (29%) than dSPNs (10%) were modulated by uncertainty (p = 0.0023, Chi-squared), suggesting a selective role for iSPNs in hesitation. Indeed, during Uncertain (**Fig. 3E**), but not Rewarded or Unrewarded (**Fig. S2D,E**) trials, there was a stronger negative spike/lick correlation in the uncertainty-responsive iSPN population (r = -0.12 ± 0.03) compared with the positive value (r = 0.007 ± 0.01), negative value (r = -0.01 ± 0.02), or certainty (r = -0.003 ± 0.03) populations. Interestingly, there were no significant spike/lick correlations in any population of dSPNs on Uncertain (**Fig. 3H**), Unrewarded (**Fig. S2F**), or Rewarded (**Fig. S2G**) trials. Consequently, a population of DMS iSPNs, but not dSPNs, is tuned to uncertain tones and covaries with anticipatory licking. These characteristics position DMS iSPNs to mediate hesitation.

**Figure 3:**
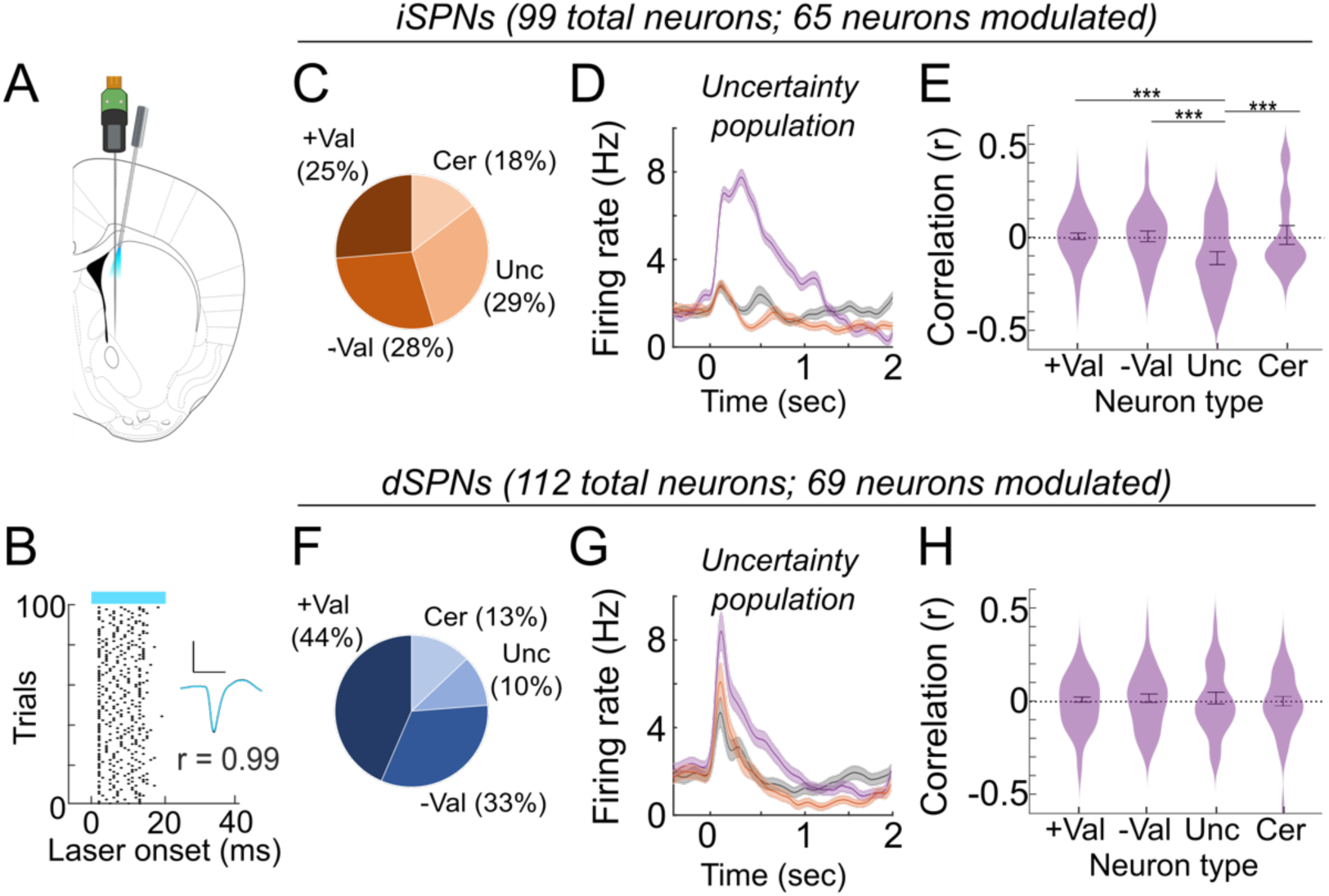
Indirect pathway SPNs are enriched in uncertainty-responsive neurons that correlate with hesitation. **A**) We optogenetically identified responses from iSPNs and dSPNs using a custom made optrode (see Methods). **B**) After each session, we stimulated neurons expressing ChR2 100 times (473mW, 20ms, 1mW) at 1Hz. ‘Tagged’ neurons were identified based on the correlation between laser-driven and spontaneous spikes and spike reliability following laser stimulation (see Methods). Example of ‘tagged’ iSPN. *Inset axes:* 400uV (y-axis) and 1ms (x-axis). **C-E**) Data from 99 identified iSPNs (4 mice, 9 sessions). **C**) Proportions of modulated neurons that fall into each of the four response types. **D**) Average PSTH from the uncertainty-responsive iSPNs. **E**) Violin plots of the distribution of spike/lick correlations across the four types of neural responses for iSPNs. **F-H**) Same as C-E but for dSPNs (112 neurons; 3 mice; 8 sessions).

### Optogenetic activation of indirect pathway neurons induces hesitation

To determine whether iSPNs play a causal role in hesitation, we transiently stimulated DMS iSPNs during the cue period of the task (**Fig. S3A-B**). Activating iSPNs during Uncertain cues significantly delayed the onset of anticipatory licking (**Fig. 4A-C**; p < 0.001, paired t-test). In contrast, stimulation during the Unrewarded (**Fig. 4D-F**) or Rewarded (**Fig. 4G-I**) cues did not change anticipatory licking, suggesting that iSPNs exclusively influence behavior during Uncertain trials. Interestingly, activating iSPNs during the delay (**Fig. 4J-L**) or outcome (**Fig. S3F**) periods of Uncertain trials did not alter licking, suggesting that the iSPNs influence hesitation rather than lick rate. iSPN activation during the delay or outcome periods of Unrewarded (**Fig. S3C,E**) or Rewarded (**Fig S3D,G**) trials did not alter licking, further supporting the selective role of iSPNs in hesitation. Importantly, there were no changes in anticipatory licking if light stimulation occurred in non-ChR2 control animals (**Fig. S3H-J**). Finally, because striatal direct and indirect pathways often play opposing roles in behavior^16–18^, we tested whether activating dSPNs would eliminate hesitation. Surprisingly, optogenetic stimulation of dSPNs during the cue, delay, or outcome periods of Unrewarded, Uncertain, or Rewarded trials never altered licking (**Fig. S4**). Consequently, contrary to traditional models of striatal function, activating DMS iSPNs, but not dSPNs, influenced hesitation.

**Figure 4:**
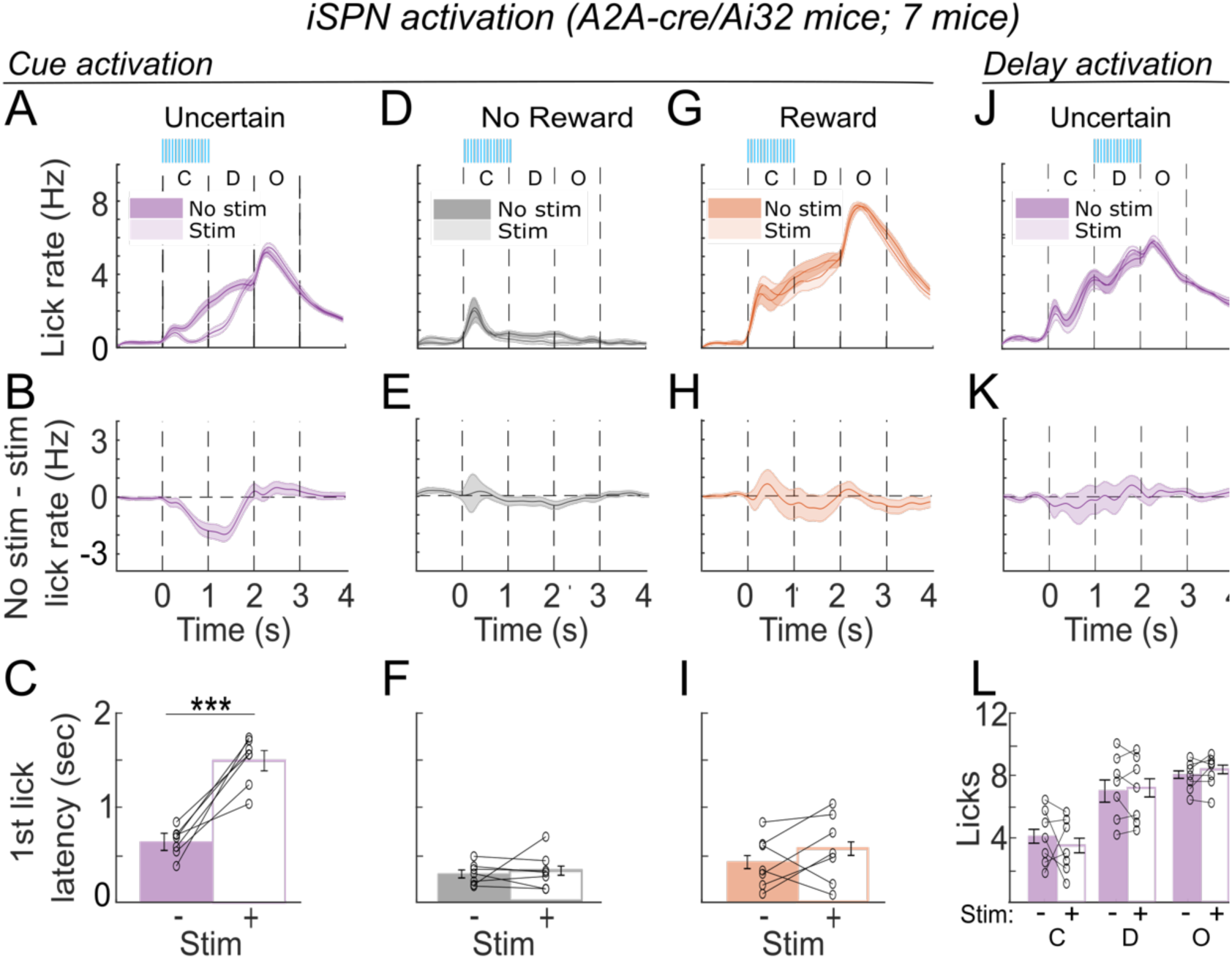
Optogenetic activation of iSPNs exaggerates hesitation. Mice which expressed ChR2 in iSPNs (A2A-cre/Ai32) were implanted bilaterally with fibers in the DMS (n=7; 3 females). We stimulated iSPNs during the cue period (473nm; 16 Hz, 10ms pulses; 2mW). **A-C**) Effect of iSPN activation during uncertain trials. **A**) Comparison of licking during stimulation (light) and control (dark). **B**) Difference in licking between stimulation and control trials. **C**) Median first lick latency for each animal during stimulation and control sessions. Data from each animal are taken from 3 stimulation sessions and 3 control sessions (see Methods). **D-F**) Same as a-c but for stimulation during unrewarded trials. **G-I**) Same as a-c but for rewarded trials. **J-K**) Comparison of licking during stimulation during the delay period of uncertain trials. **J**) Comparison between stimulation and control trials. **K**) Difference in licking between stimulation and control trials. **L**) Average number of licks during the cue, delay, and outcome period for each animal during stimulation and control trials. *** indicates p < 0.001 in paired t-test.

We next tested whether other striatal subregions played similar roles in hesitation. We focused on the ventrolateral striatum (VLS) as it has been shown to regulate licking^19–21^ (**Fig. S5A-B**). Optogenetic activation of VLS iSPNs during either the cue (**Fig. S5C-E**) or outcome (**Fig. S5F-H**) periods of any trial type reduced licking, suggesting that the VLS controls lick rate rather than hesitation. Conversely, activating VLS dSPNs during the cue (**Fig. S5I-K**) or outcome (**Fig. S5L-N**) periods increased anticipatory licking across all trial types. Therefore, while VLS iSPNs and dSPNs bidirectionally control lick rate, only DMS iSPNs regulate hesitation.

Finally, we asked whether the magnitude of hesitation varies with the level of uncertainty. We trained a separate cohort of mice on a variation of the task in which mice were rewarded on 66% of Uncertain trials instead of 50% (**Fig. S6A**). Interestingly, while mice continued to hesitate on the 66% Uncertain trials, decreasing the amount of uncertainty also decreased hesitation. Mice began licking approximately 500ms after the onset of the 66% Uncertain tone, compared to approximately 750ms on the 50% Uncertain tone (**Fig. S6B,C**). Similar to the results of the initial task, iSPN activation on 66% Uncertain trials, but not other trial types, exaggerated hesitation (**Fig. S6C-H**). Consequently, hesitation correlates with the level of uncertainty, and DMS iSPNs control hesitation across multiple levels of uncertainty.

### Optogenetic inhibition of indirect pathway spiny projection neurons (iSPNs) reduces hesitation

Finally, we silenced DMS iSPNs during each of the three auditory cues to test whether iSPNs are necessary for hesitation. Inhibiting iSPNs during the Uncertain cue eliminated hesitation (**Fig. 5A-C**; p < 0.001, paired t-test), whereas inhibiting iSPNs during Unrewarded (**Fig. 5D-F**) or Rewarded (**Fig. 5G-I**) cues did not affect licking. Additionally, inhibiting iSPNs during the delay period did not alter the rate or timing of anticipatory licking (**Fig. 5J-L; Fig. S7A,B**). Finally, inhibiting dSPNs during any of the three cues did not alter anticipatory licking (**Fig. S7C-D**). Taken together, our electrophysiology and optogenetic experiments indicate that DMS iSPNs control hesitation.

**Figure 5:**
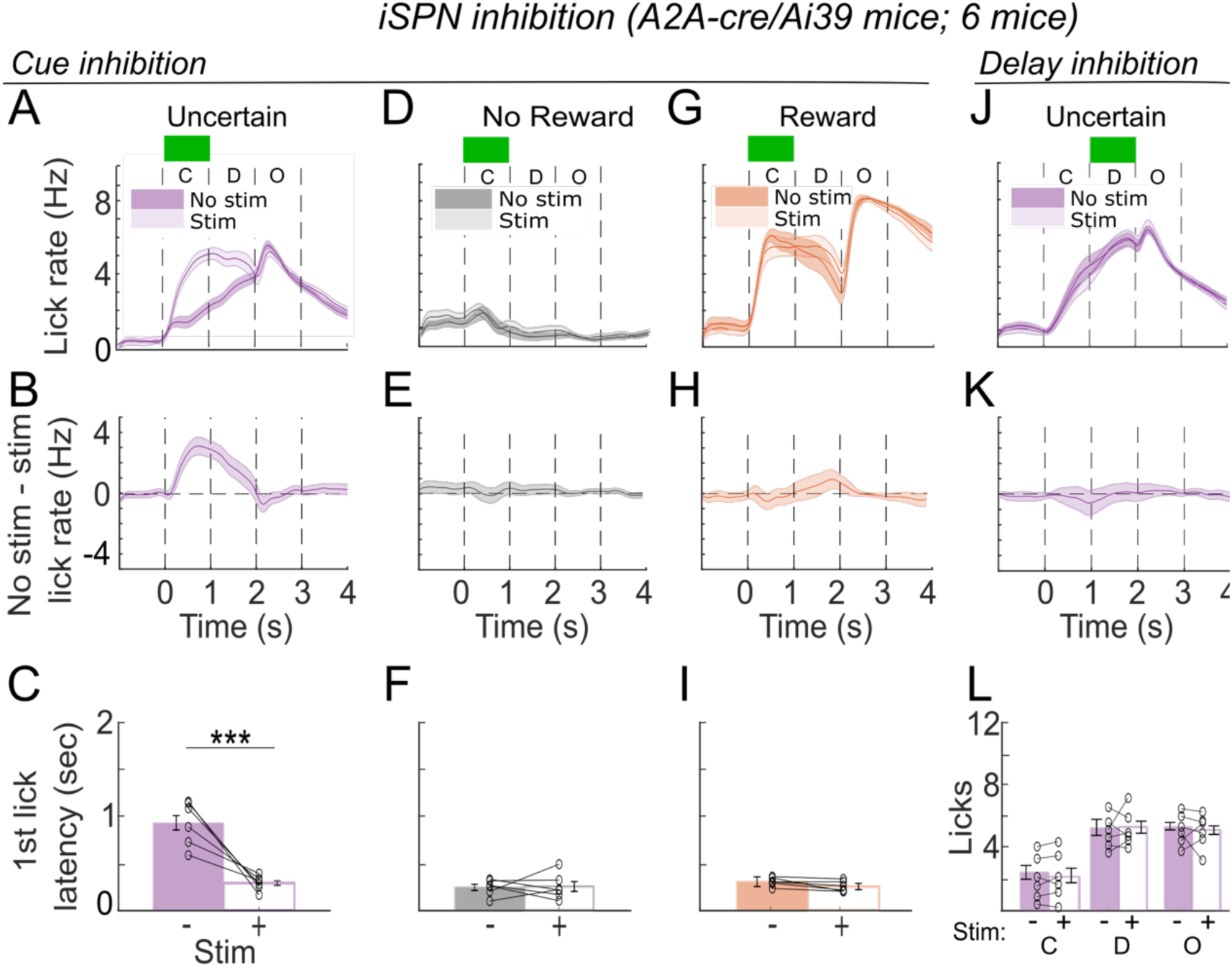
Optogenetic inhibition of iSPNs eliminates hesitation. Mice which expressed halorhodopsin in iSPNs (A2A-cre/Ai39) were implanted bilaterally with fibers in the DMS (n=6; 2 females). We inhibited iSPNs during the cue period (532nm; 1sec; 5mW). **A-C**) Effect of iSPN inhibition during uncertain trials. **A**) Comparison of licking during inhibition (light) and control (dark) trials. **B**) Difference in licking between inhibition and control trials. **C**) Median first lick latency for each animal during inhibition and control sessions. **D-F**) Same as a-c but for inhibition during unrewarded trials. **G-I**) Same as a-c but for rewarded trials. **J-K**) Comparison of licking during inhibition during the delay period of uncertain trials. **J**) Comparison between inhibition and control trials. **K**) Difference in licking between inhibition and control trials. **L**) Average number of licks during the cue, delay, and outcome period for each animal during inhibition and control trials. *** indicates p < 0.001 in paired t-test.

## Discussion

Hesitation is a critical cognitive function that allows for deliberation and often leads to more accurate decisions^3,22^. Despite the widespread prominence of hesitation in both health and disease, little is known about its underlying neural mechanisms. Here, using an auditory association task, we present the first behavioral paradigm that allows the reliable study of hesitation in rodents. We show that activity in DMS iSPNs specifically correlates with hesitation, and that bidirectional modulation of iSPN, but not dSPN, activity alters hesitation. At first glance, our findings that the indirect pathway circuit mediates hesitation appear to merely corroborate its proposed role in general movement suppression^17^. Yet hesitation is more than simply suppressing movement: it is an uncertainty-dependent pause before further action. These results indicate that hesitation is mediated by a separate basal ganglia circuit compared to cue-induced stopping and suggest that an organism’s specific behavioral needs dictates the mechanisms that will ultimately inhibit action^6,10,16,23^.

For hesitation to occur, the brain must rapidly identify uncertain contexts that warrant a pause in behavior. Frontal areas, the anterior cingulate cortex (ACC) in particular, regulate a variety of uncertainty-based behaviors such as information seeking and probabilistic reversal learning^27–29^ and preferentially target this dorsomedial portion of striatum^25,26^, making them promising candidates to also mediate hesitation. Indeed, an ACC neurons signal the uncertainty of cue-reward associations^29^ and the volatility of the reward environment^30,31^. DMS neurons also receive some inputs from motoric cortex^32^, opening the question as to whether the decision to hesitate is inherited from a single upstream area or whether DMS iSPNs integrate converging uncertainty signals from areas such as the ACC and movement signals from motor cortex.

We also recognize that although hesitation may be useful for providing additional time to deliberate during difficult decisions^33^, it is unclear why mice hesitate during this task when there is no decision to be made. While additional time may be of consequence in some scenarios, this needn’t be the case. In a two-choice task, when monkeys were observed hesitating before performing an action in an uncertain context, the represented motor plan observed in premotor or motor cortex during this time did not change^34^. These surprising results indicate that hesitation may simply be a cognitively driven pause in action execution, not the replanning or vacillation between two options. More broadly, hesitation may be unavoidable in many circumstances, even if it does not directly serve to improve performance. Rather than being a consequence of computationally intensive decision-making systems, the opposite may be true: the presence of hesitation in a variety of uncertain scenarios may have allowed organisms the necessary time to engage in and evolve complex decision-making processes.

Our findings are also notable given the influential ‘race models’ of action inhibition, based on data from stop-signal tasks^4–6,10^, that hypothesize that stopping is controlled by competing Go and Stop processes mediated by the basal ganglia direct and hyperdirect pathways, respectively^4,5^. While our data do not conclusively rule out a ‘race’ mechanism, the data suggest a very different process that does not involve competition between opponent operators. Because dSPNs are not modulated by hesitation and manipulating DMS dSPN activity does not impact hesitation, dSPNs cannot mediate a Go process in this context. Likewise, while we cannot rule out a modulating influence of the hyperdirect pathway in a Stop process, we demonstrate that iSPN activity is sufficient and necessary for hesitation.

Consequently, we propose that multiple different basal ganglia subcircuits can suppress actions depending on the specific behavioral objective, potentially all utilizing the same mechanisms to orchestrate the desired pause (e.g. SNr dynamics). Thus, while existing conceptual frameworks are invaluable, they are insufficient to capture more cognitively demanding mechanisms for stopping such as during hesitation.

## Acknowledgments

We thank Drs. A. Gittis, R. Peixoto, B. Chamberlain, and members from the Yttri and Ahmari labs for thoughtful feedback on the manuscript.

## Funding

CRCNS R01 – DA059993 (EAY)

CRCNS R01 – DA053014 (EAY)

US-Israel Binational Science Foundation

(EAY) NIMH R01MH119837 (SEA)

NINDS R01NS125141 (SEA)

## Author contributions

Conceptualization: MAG, EAY

Methodology: MAG, EAY

Investigation: MAG, EAY

Visualization: MAG, EAY

Funding acquisition: MAG, SEA, EAY

Project administration: MAG, EAY

Supervision: MAG, SEA, EAY

Writing – original draft: MAG, SEA, EAY

Writing – review & editing: MAG, SEA, EAY

## Data and materials availability

All data and code from the main text and the supplementary materials will be posted on Github upon publication.

## Supplemental Materials

### Methods

#### Mice

Experiments were conducted in accordance with the guidelines from the National Institutes of Health and with approval from Carnegie Mellon University Institutional Animal Care and Use Committee. Adult male and female mice (>8 weeks old) were used for experiments. All mice were maintained on a reverse circadian 12-h light/12-h dark cycle with food provided ad libitum. Drd1-cre and A2A-cre mice were used to selectively target D1+ or D2+ neurons, respectively. To activate or silence either D1+ or D2+ mice, we crossed these transgenic lines with ChR2 reporter mice (Ai32) or halorhodopsin reporter mice (Ai39), respectively.

#### Surgery

Mice were anesthetized using 4% isoflurane mixed with oxygen and maintained on 1-2% isoflurane for the duration of surgery. Mice were placed on a small-animal stereotactic instrument and secured using ear and bite bars. Hair was removed from the dorsal surface of the head with hair clippers and the incision area was scrubbed with a betadine solution. A large incision was then made exposing the dorsal portion of the skull. AP and ML measurements were made relative to an interpolated bregma; DV measurements were made relative to dura. For optogenetic experiments, optical fibers were implanted bilaterally into the DMS (AP: 0.7mm; ML: 1.5mm; DV: 2.5mm) or VLS (AP: 0.7mm; ML: 2.5mm; DV: 3.5mm) and secured using dental cement. Custom 3D printed headcaps (https://github.com/YttriLab/Joystick/tree/master/Mouse%20Shuttle%20Parts) were also secured onto the skull using dental cement for head-fixation. For electrophysiology experiments, we performed two surgeries. In the first surgery, the skull over the DMS (AP: 0.7mm; ML: 1.5mm) was marked, and a headcap was secured onto the skull. After we trained the animal on the task, we implanted a skull screw attached to a silver wire over the cerebellum and made a craniotomy over the previously marked DMS coordinates.

#### Auditory association task

Prior to training, mice were handled and water restricted to 1ml daily for one week. Once training started, mice were headfixed and placed in front of a lick spout. Deflections in the lick spout caused by licking were measured using a piezoelectric sensor (LDT1-028K Piezo Sensor; Newark). Individual licks were detected post-hoc using custom scripts written in Matlab (Mathworks). The task was controlled using custom scripts run through Arduino microcontroller. Three tones (70dB; 12kHz pure tone, 3kHz pure tone; and white noise) were generated using a stereo 20W class D audio amplifier (MAX9744; Adafruit). Tone-reward associations (100% reward, 50% reward, or 0% reward) were counterbalanced across animals. On each trial, one tone was presented for 1sec followed by a 1sec delay period (unless otherwise specified). After the delay period, the trial outcome was signaled by either a 4uL drop of sweetened water (3mM acesulfame potassium) or a 200msec house light. Training consisted of 3 stages. In the first stage, each tone was presented in blocks of 10 trials. Each session consisted of 5 blocks for each tone for a total of 150 trials. The first stage was further divided into substages in which the length of the delay period varied. In the first substage, the delay period was 250ms. The delay period increased in 250ms intervals up to 1sec. In the second stage, each tone was presented in blocks of 4 trials. In the final stage, tones were presented pseudorandomly in blocks of 30 trials for a total of 300 trials. Each block of 30 trials consisted of 10 rewarded trials, 10 unrewarded trials, and 10 uncertain trails (5 rewarded and 5 unrewarded). Animals advanced to the next stage or substage of training once they showed anticipatory licking during the cue and delay period that correlated with the value of the tone for two consecutive days. Training lasted between 16 and 22 days.

#### Optogenetics

For all implants, 3.0mm optical fibers were glued inside 1.25mm ceramic ferrules. All fibers were calibrated to achieve the desired laser output of 2mW. Excitation of dSPNs or iSPNs was performed using pulsed 2mW 473nm light (16Hz, 10ms pulses) for 1 sec from a LED light source (Pritzmatix). Stimulation parameters were chosen as they have been previously shown to induce physiological increases in SPN firing rate without altering locomotion^16^. Inhibition of dSPNs or iSPNs was performed using constant 5mW 532nm light from a laser (Opto Engine, Midvale Utah USA). Each week, we manipulated neural activity during a distinct 1sec period (**Fig. S3B**). For example, one week consisted of optogenetically manipulating activity during the cue period of uncertain trials. Three days per week (either M,W,F or T,Th,Sat depending on animal) the laser would be active. On the other three days, the animal would still be connected to the light source but the laser would be inactive. On days when the laser was active, no stimulation would occur during the first and last 20 uncertain trials. Stimulation only occurred between the 21^st^ and 79^th^ presentation of the uncertain trials. This experimental design allowed us to control for within and across day variation in anticipatory licking.

#### Neuropixels recordings

Electrophysiology recordings were made with Neuropixels^11^ Phase 3A probes connected to a micromanipulator (Scientifica) through a metal rod. Raw data within the action potential band (band-pass filtered from 0.3-10kHz, 30kHz sampling) were collected using SpikeGLX software (https://github.com/billkarsh/SpikeGLX). Probes were coated with DiI fluorescent dye to histologically confirm placement in the DMS. Data were spike-sorted using Kilosort 3.0 (https://github.com/MouseLand/Kilosort) and manually curated using Phy (https://github.com/cortex-lab/phy).

#### Single-unit analysis and cell-type classification

Neurons were considered to be in the DMS if they were within 1.0mm from the corpus-collosum. The corpus-collosum was clearly identifiable as an absence of units after processing in kilosort. High-quality DMS single units were defined by the following criteria^35^: 1) waveform amplitude of more than 50 uV; 2) greater than 300 spikes; 3) estimated false-positve rate less than 10%; 4) waveform trough that preceded the peak to eliminate axonal spikes. DMS single units were next classified into one of four striatal cell types^35^: spiny projection neurons (SPNs), fast-spiking interneurons (FSIs), tonically active neurons (TANs), or unidentified interneurons (UINs) using previously published criteria. Spike trough-to-peak duration was first used to separate SPNs and TANs (> 400us) from FSIs and UINs (<= 400us). Post-spike suppression (PSS) was then used to differentiate SPNs (PSS <= 40ms) from TANs (PSS > 40ms). FSIs and UINs were differentiated by calculating the proportion of inter-spike intervals (ISIs) that were greater than 2s. Neurons with a proportion greater than 10% were UINs, while neurons with a proportion less than 10% were FSIs.

#### Identification of light-responsive neurons

To perform cell-type specific recording from dSPNs or iSPNs, an optical fiber was attached to the side of the Neuropixels probe at a 15° angle. The tip of the fiber was 250um from the probe. At the end of a recording session, 100 laser pulses (473mW, 20ms, 1mW) were delivered at 1Hz to identify light-responsive neurons. Neurons were considered light-responsive if two criteria were met as previously described^14,36^. First, the correlation between laser-driven and spontaneous spike waveforms must be >=0.95. Second, the first spike latency in the first 10ms window after the onset of laser stimulation must be significantly lower than the first spike latency in 100 randomly chosen 10ms windows in between laser pulses. (log-rank test, p<0.01).

#### Neural data analysis

We first identified DMS SPNs that were differentially responsive to the three tones (ANOVA, p<0.05) in a 740ms time window that started at tone onset. We chose to analyze responses in the 740ms window because it is the median first lick latency during uncertain trials, and we were primarily interested in uncertainty-responsive neurons that could influence hesitation. Neurons that were differentially responsive to the tones, as identified by a significant ANOVA, were further subdivided into positive value-, negative value-, uncertainty-, or certainty-responsive neurons based on their specific responses to the tones. The following definitions are based on t-tests between firing rates and are exhaustive such that all responsive neurons fell into exactly one of the four groups. Neurons were ‘Positive-value-responsive’ if the firing rates differed between the 3 tones as follows: (0 < 100) & (0 <= 50) & (50 <= 100). Analogously, neurons were ‘negative-value-responsive’ if the firing rates varied such that: (100 < 0) & (50 <=0) & (100 <= 50). Neurons were uncertainty-responsive if: (0 < 50) & (100 < 50). Finally, neurons were certainty-responsive if: (50 < 0) & (50 < 100). To determine the covariation between spikes and licks, we took the correlation between the number of spikes and licks in a 740ms time window starting at tone onset.

## Supplemental Figures

**Figure S1:**
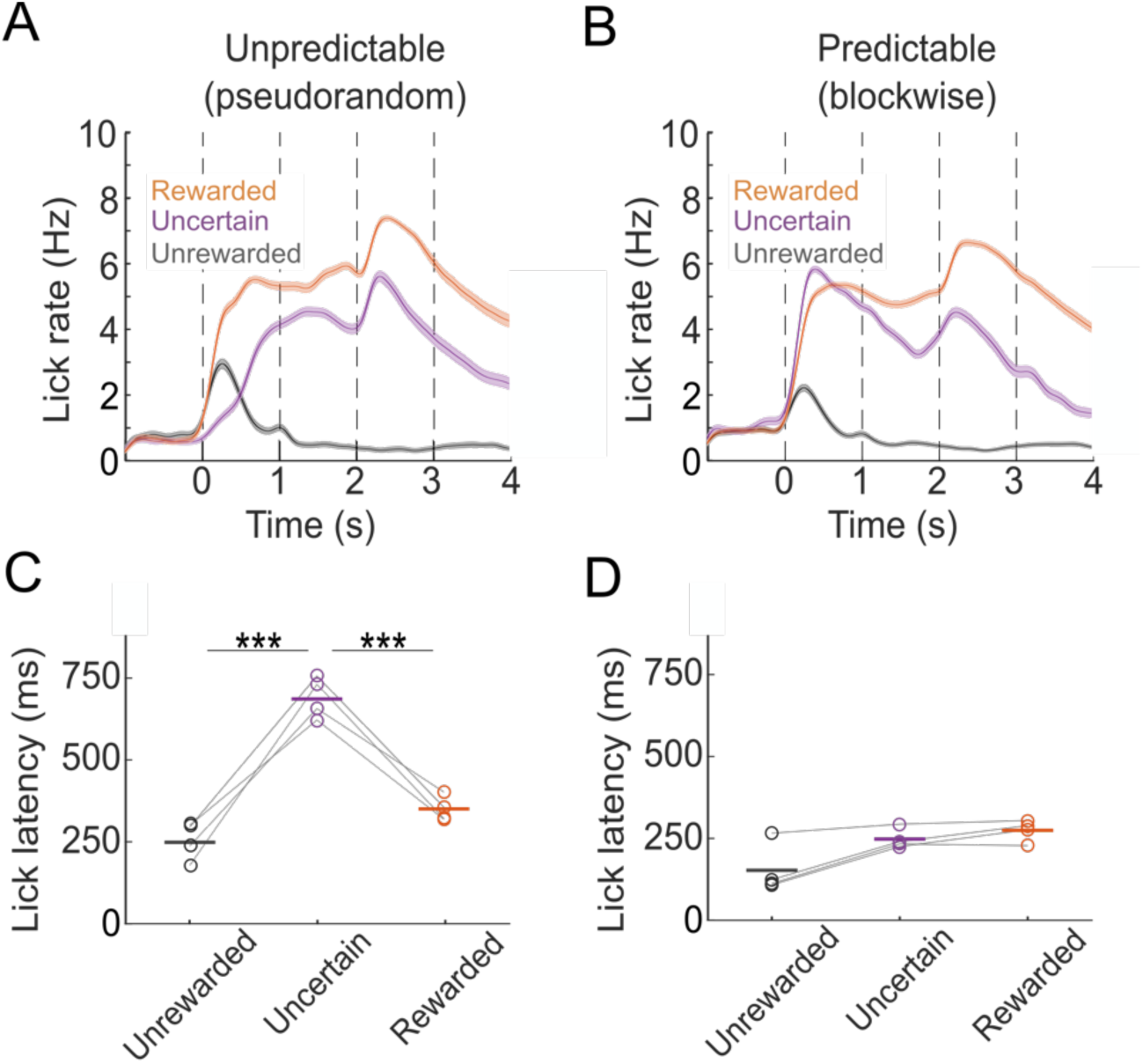
Mice only hesitate if tones are unpredictable. A subset of mice was trained on two different versions of the task. Mice first performed a version of the task when tones were presented pseudorandomly and thus unpredictable. They then performed a version of the task with tones that were presented in blocks of 10 and thus predictable. **A-B**) Anticipatory licking correlated with tone value for unpredictable (**A**) and predictable tones (**B**). **C-D**) First lick latency for all tones when tones were unpredictable (**C**) or predictable (**D**). Mice hesitated when tones were unpredictable but not predictable.

**Figure S2:**
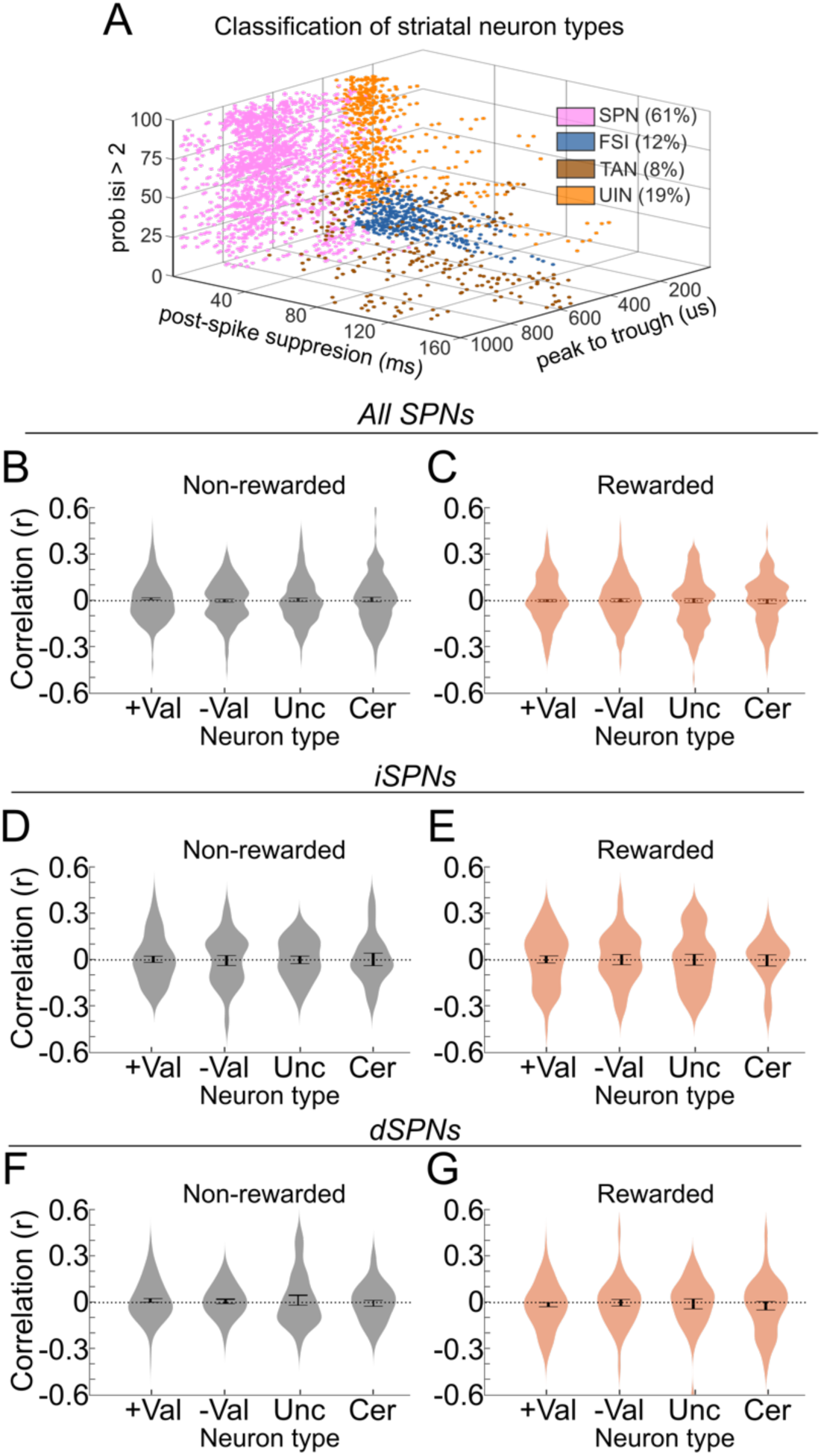
Additional electrophysiology analyses. **A**) Classification of 1717 DMS neurons from 9 animals (24 sessions). Neurons were classified into SPNs (pink), FSIs (blue), TANs (brown), or UINs (orange) based on waveform peak-to-trough duration, post-spike suppression, or percent of inter-spike-intervals greater than 2 sec (see methods). **B-C**) Distribution of spike-lick correlations across 4 classes of neurons (+Val, -Val, Unc, Cer) for unrewarded trials (**B**) and rewarded trials (**C**). Data taken from all recorded SPNs. Uncertain trials displayed in Fig 2h. **D-E**) Same as b-c but for identified iSPNs. Uncertain trials displayed in Fig 3e. **F-G**) Same as b-c but for identified dSPNs. Uncertain trials displayed in Fig. 3h. For B-G, no average correlation was significantly different from zero.

**Figure S3:**
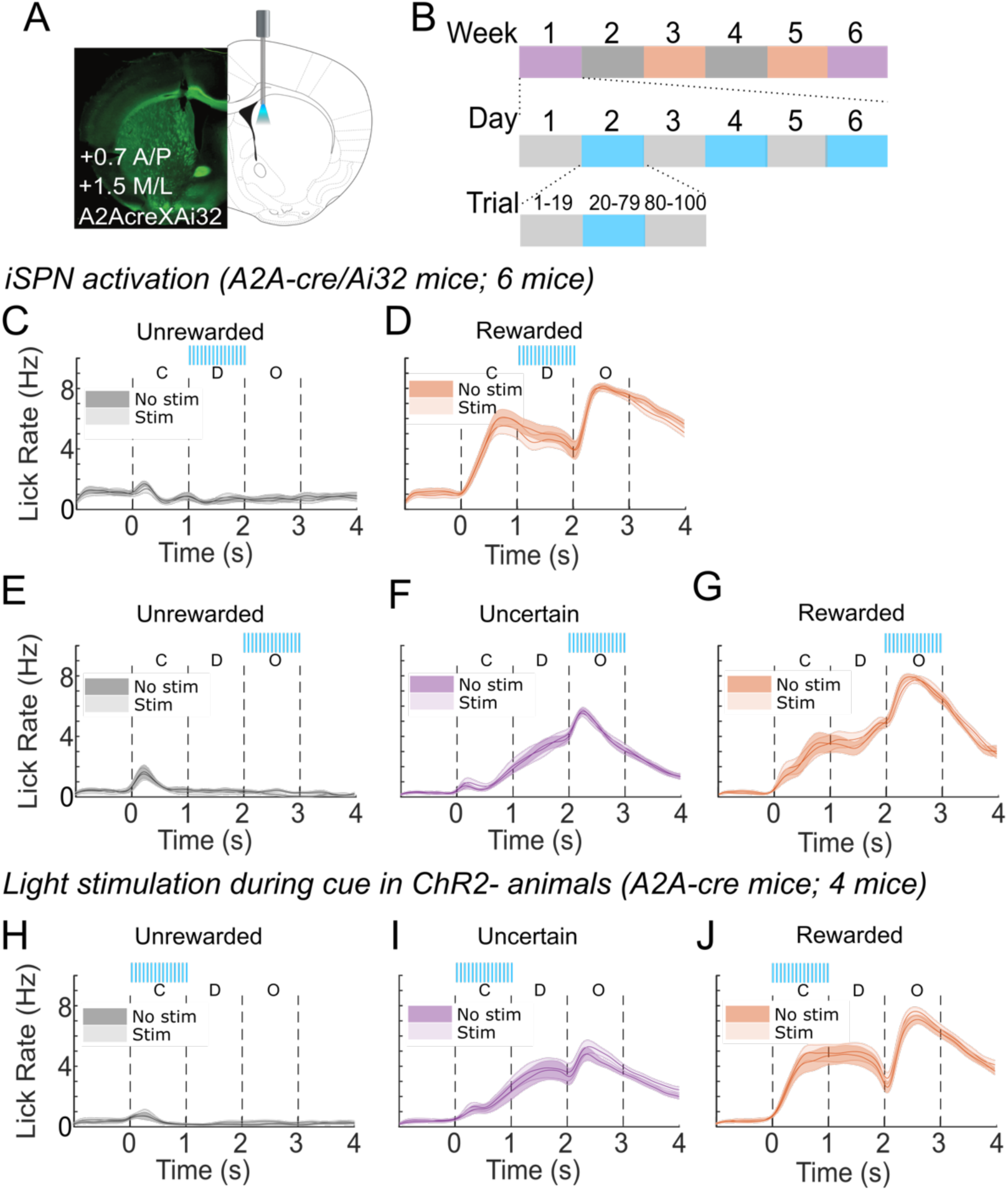
Additional optogenetic activation data. **A**) Histology of fiber placement in the DMS of A2AcreXAi32 mouse. **B**) Overview of optogenetic activation experiment. Each week only a single trial type was activated on alternating days (see methods). Each day of stimulation (blue), the first and last 20 trials were unstimulated. **C-D**) Optogenetic activation of DMS iSPNs (6 mice) during the delay period of unrewarded (**C**) or rewarded (**D**) trials. Activation during delay period of uncertain trials displayed in Fig. 4j-l**. E-G**) Optogenetic activation of DMS iSPNs during the outcome period of unrewarded (**E**), uncertain (**F**), or rewarded (**G**) trials. **H-J**) Light illumination of DMS in animals (A2A-cre; 4 mice) that do not express ChR2 during unrewarded (**H**), uncertain (**I**), or rewarded (**J**) trials.

**Figure S4:**
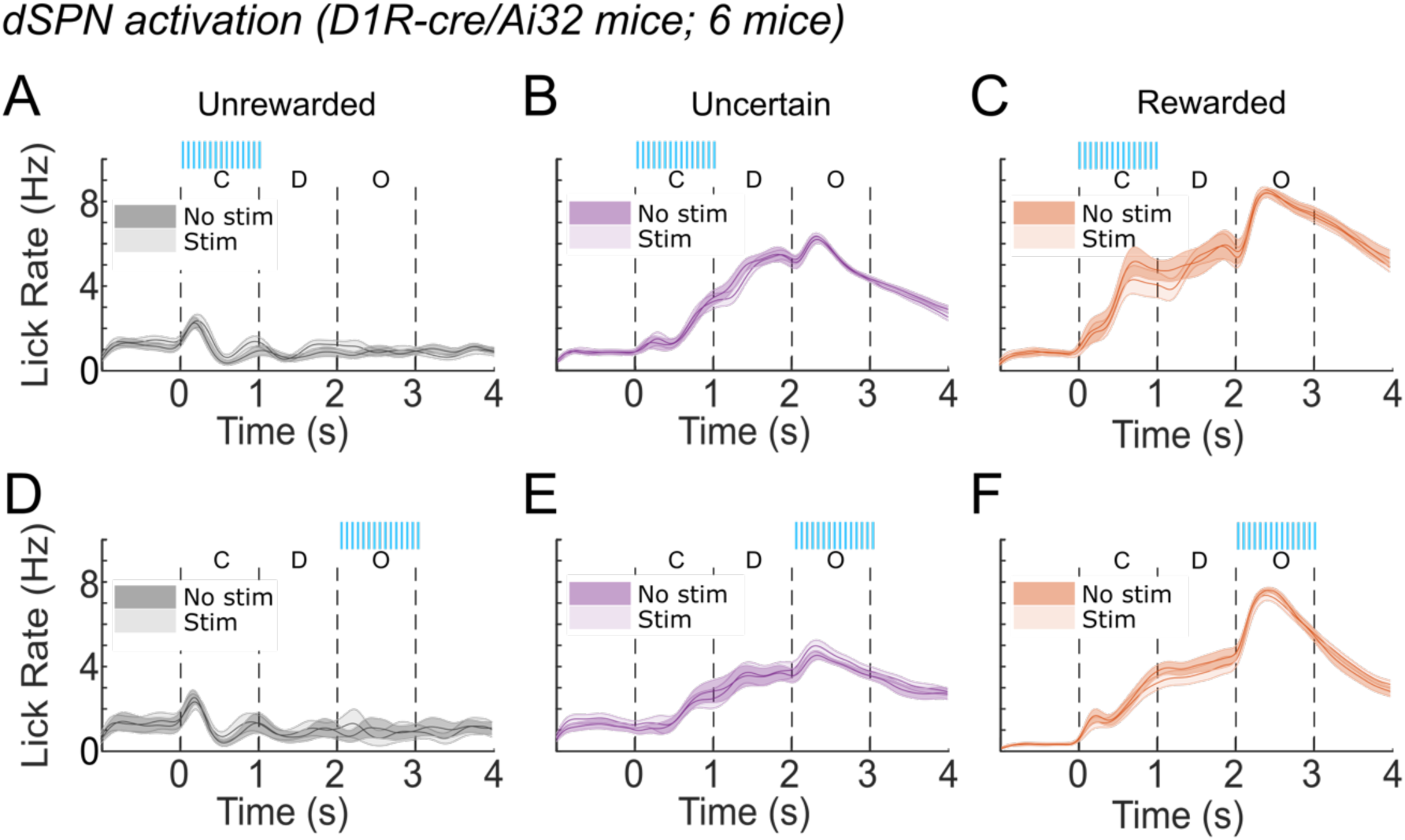
Optogenetic activation of dSPNs does not influence licking. Optogenetic activation of DMS iSPNs (6 mice) during the auditory association task. **A-C)** Activation of dSPNs during the delay period of unrewarded (**A**), uncertain **(B**), or rewarded (**C**) trials does not influence licking. **D-F)** Same as a-c but for optogenetic stimulation of dSPNs during the outcome period.

**Figure S5:**
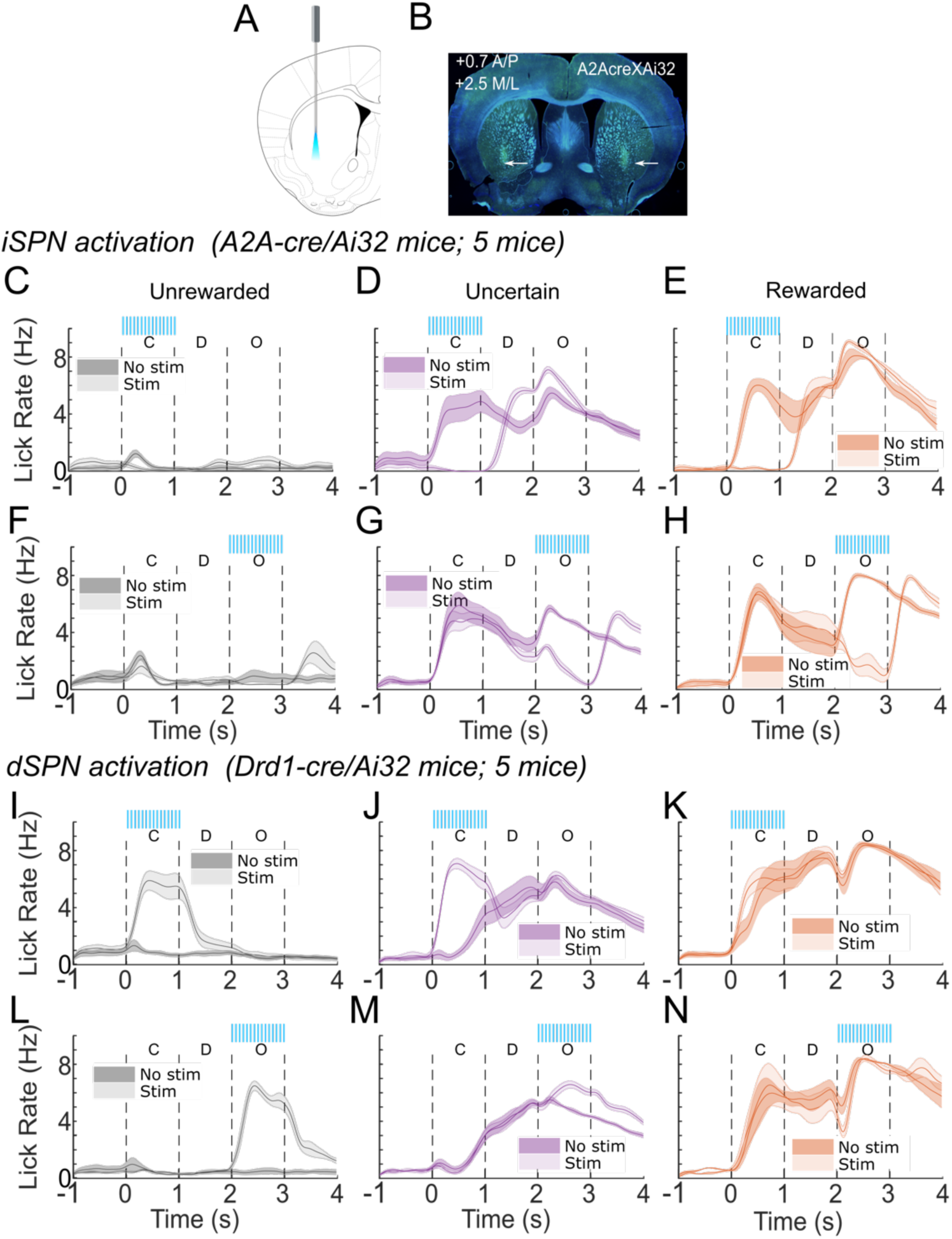
Optogentic activation of VLS iSPNs and dSPNs. **A**) Mice were implanted with bilateral optical fibers in the VLS (AP: 0.7mm; ML: 2.5mm; DV: 3.5mm). **B**) Histological confirmation of optical fibers in the VLS. **C-E**) Optogenetic activation of iSPNs during the cue period of unrewarded (**C**), uncertain (**D**), or rewarded (**E**) trials all decreased anticipatory licking. **F-H**) Same as C-E but with activation of iSPNs during the outcome period. **I-K)** Activation of dSPNs during the cue period of unrewarded (**I**), uncertain (**J**), or rewarded (**K**) trials in the VLS increased anticipatory licking regardless of cue type. **L-N**) Same as i-k but with activation of dSPNs during the outcome period.

**Figure S6:**
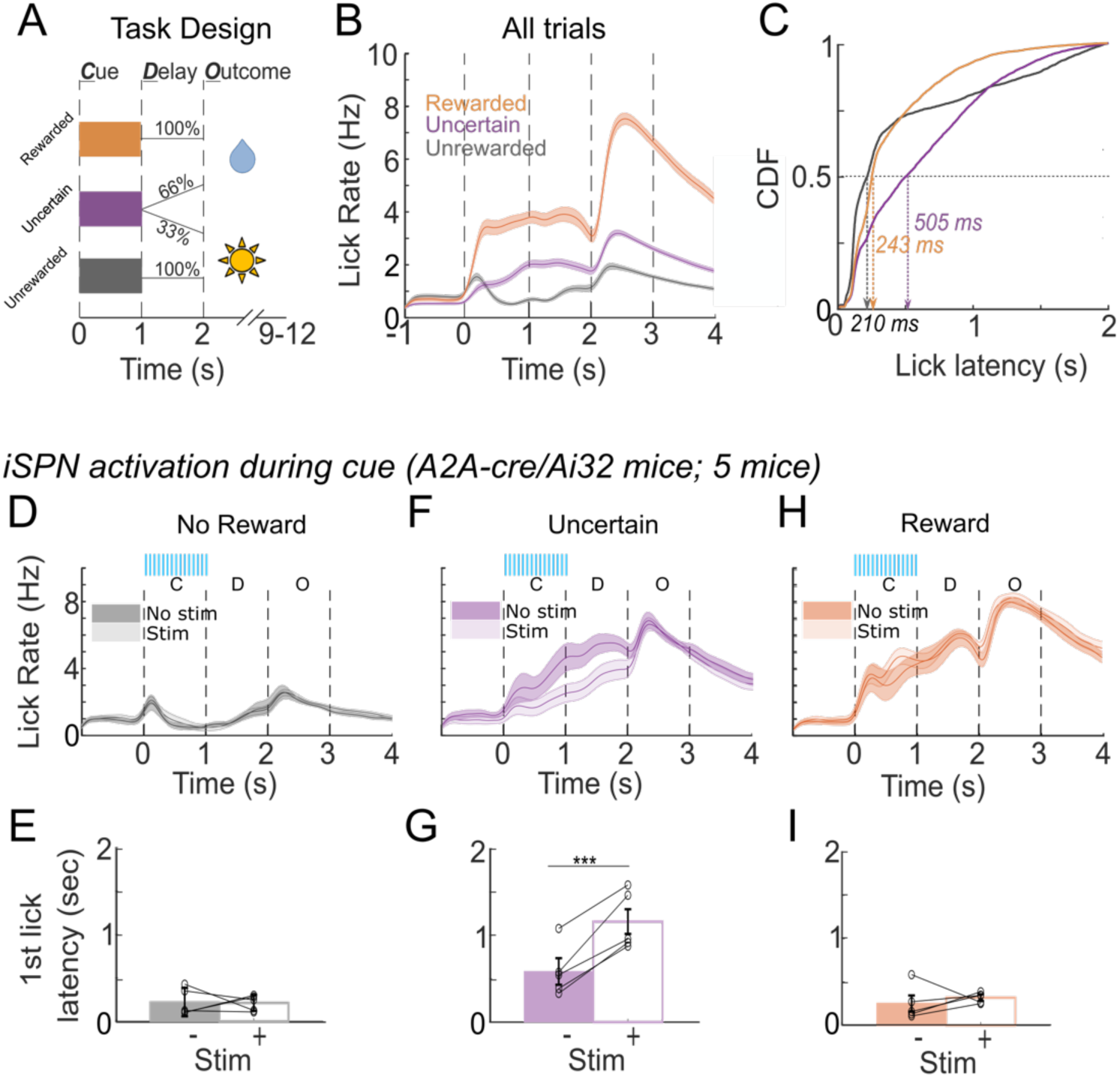
DMS iSPNs control hesitation across multiple levels of uncertainty. **A**) Overview of alternative auditory conditioning task. Similar to the original task, one tone is never rewarded (gray) and a second tone is always rewarded (orange). A third tone is rewarded on 66% of trials, instead of 50% of trials as in the original task to decrease the amount of uncertainty. **B**) Similar to the original task, mice lick in anticipation to the reward in a manner that reflects tone value. **C**) First lick latency during the uncertain trials is longer compared with unrewarded or rewarded trials. **D-E**) Anticipatory licking (**D**) or first lick latency (**E**) does not change after optogenetic activation during the cue period of unrewarded trials. **F-G**) Same as d-e but with stimulation during the uncertain trials. Activation of iSPNs during the uncertain trials decreases anticipatory licking (**F**) and increases first lick latency (**G**; p< 0.001, paired t-test). **H-I**) Same as d-e, but with optogenetic stimulation during rewarded trials.

**Figure S7:**
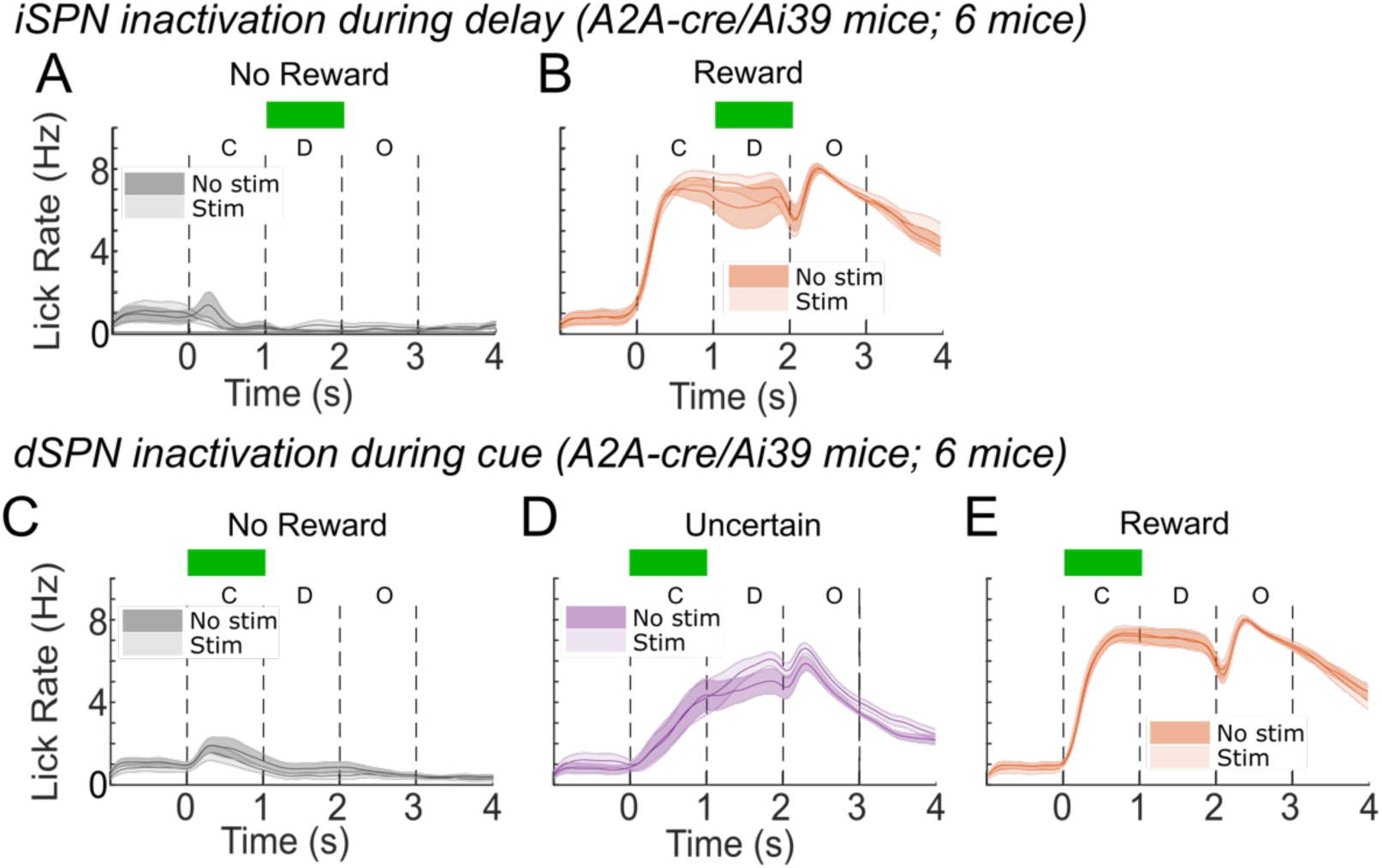
Optogenetic inhibition of DMS dSPNs does not alter change hesitation. **A,B**) Inhibition of DMS iSPNs (6 mice) during the delay period of unrewarded (**A**) or rewarded trials (**B**). Inhibition of iSPNs during the delay period of uncertain trials is displayed in Fig. 5a-c. **C-E**) Inhibition of DMS dSPNs (6 mice) during the cue period of unrewarded (**C**), uncertain (**D**), or rewarded (**E**) trials does not change anticipatory licking.

## References

1. Merriam-Webster. Hesitate. Merriam-Webster2024.

2. Redish AD. Vicarious trial and error. Nat Rev Neurosci. Mar 2016;17(3):147-59. doi:10.1038/nrn.2015.30

3. Frank MJ, Samanta J, Moustafa AA, Sherman SJ. Hold your horses: impulsivity, deep brain stimulation, and medication in parkinsonism. Science. Nov 23 2007;318(5854):1309–12. doi:10.1126/science.1146157

4. Aron AR. From reactive to proactive and selective control: developing a richer model for stopping inappropriate responses. Biol Psychiatry. Jun 15 2011;69(12):e55–68. doi:10.1016/j.biopsych.2010.07.024

5. Aron AR, Behrens TE, Smith S, Frank MJ, Poldrack RA. Triangulating a cognitive control network using diffusion-weighted magnetic resonance imaging (MRI) and functional MRI. J Neurosci. Apr 04 2007;27(14):3743–52. doi:10.1523/JNEUROSCI.0519-07.2007

6. Schmidt R, Berke JD. A Pause-then-Cancel model of stopping: evidence from basal ganglia neurophysiology. Philos Trans R Soc Lond B Biol Sci. Apr 19 2017;372(1718)doi:10.1098/rstb.2016.0202

7. Schall JD, Palmeri TJ, Logan GD. Models of inhibitory control. Philos Trans R Soc Lond B Biol Sci. Apr 19 2017;372(1718)doi:10.1098/rstb.2016.0193

8. Logan GD, Cowan WB, Davis KA. On the ability to inhibit simple and choice reaction time responses: a model and a method. J Exp Psychol Hum Percept Perform. Apr 1984;10(2):276–91. doi:10.1037//0096-1523.10.2.276

9. Chen W, de Hemptinne C, Miller AM, et al. Prefrontal-Subthalamic Hyperdirect Pathway Modulates Movement Inhibition in Humans. Neuron. May 20 2020;106(4):579–588.e3. doi:10.1016/j.neuron.2020.02.012

10. Schmidt R, Leventhal DK, Mallet N, Chen F, Berke JD. Canceling actions involves a race between basal ganglia pathways. Nat Neurosci. Aug 2013;16(8):1118–24. doi:10.1038/nn.3456

11. Jun JJ, Steinmetz NA, Siegle JH, et al. Fully integrated silicon probes for high-density recording of neural activity. Nature. Nov 08 2017;551(7679):232–236. doi:10.1038/nature24636

12. Bolkan SS, Stone IR, Pinto L, et al. Opponent control of behavior by dorsomedial striatal pathways depends on task demands and internal state. Nat Neurosci. 03 2022;25(3):345–357. doi:10.1038/s41593-022-01021-9

13. Cox J, Witten IB. Striatal circuits for reward learning and decision-making. Nat Rev Neurosci. 08 2019;20(8):482–494. doi:10.1038/s41583-019-0189-2

14. Shin JH, Kim D, Jung MW. Differential coding of reward and movement information in the dorsomedial striatal direct and indirect pathways. Nat Commun. Jan 26 2018;9(1):404. doi:10.1038/s41467-017-02817-1

15. White JK, Monosov IE. Neurons in the primate dorsal striatum signal the uncertainty of object-reward associations. Nat Commun. Sep 14 2016;7:12735. doi:10.1038/ncomms12735

16. Yttri EA, Dudman JT. Opponent and bidirectional control of movement velocity in the basal ganglia. Nature. 05 19 2016;533(7603):402–6. doi:10.1038/nature17639

17. Kravitz AV, Freeze BS, Parker PR, et al. Regulation of parkinsonian motor behaviours by optogenetic control of basal ganglia circuitry. Nature. Jul 29 2010;466(7306):622–6. doi:10.1038/nature09159

18. Kravitz AV, Tye LD, Kreitzer AC. Distinct roles for direct and indirect pathway striatal neurons in reinforcement. Nat Neurosci. Jun 2012;15(6):816–8. doi:10.1038/nn.3100

19. Lee J, Sabatini BL. Striatal indirect pathway mediates exploration via collicular competition. Nature. Nov 2021;599(7886):645–649. doi:10.1038/s41586-021-04055-4

20. Bakhurin KI, Li X, Friedman AD, et al. Opponent regulation of action performance and timing by striatonigral and striatopallidal pathways. Elife. Apr 23 2020;9doi:10.7554/eLife.54831

21. Chen Z, Zhang ZY, Zhang W, et al. Direct and indirect pathway neurons in ventrolateral striatum differentially regulate licking movement and nigral responses. Cell Rep. Oct 19 2021;37(3):109847. doi:10.1016/j.celrep.2021.109847

22. Bogacz R, Wagenmakers EJ, Forstmann BU, Nieuwenhuis S. The neural basis of the speed-accuracy tradeoff. Trends Neurosci. Jan 2010;33(1):10–6. doi:10.1016/j.tins.2009.09.002

23. Freeze BS, Kravitz AV, Hammack N, Berke JD, Kreitzer AC. Control of basal ganglia output by direct and indirect pathway projection neurons. J Neurosci. Nov 20 2013;33(47):18531–9. doi:10.1523/JNEUROSCI.1278-13.2013

24. Hunnicutt BJ, Jongbloets BC, Birdsong WT, Gertz KJ, Zhong H, Mao T. A comprehensive excitatory input map of the striatum reveals novel functional organization. Elife. Nov 28 2016;5doi:10.7554/eLife.19103

25. Hintiryan H, Foster NN, Bowman I, et al. The mouse cortico-striatal projectome. Nat Neurosci. Aug 2016;19(8):1100–14. doi:10.1038/nn.4332

26. Monosov IE. Anterior cingulate is a source of valence-specific information about value and uncertainty. Nat Commun. Jul 26 2017;8(1):134. doi:10.1038/s41467-017-00072-y

27. Stolyarova A, Rakhshan M, Hart EE, et al. Contributions of anterior cingulate cortex and basolateral amygdala to decision confidence and learning under uncertainty. Nat Commun. Oct 17 2019;10(1):4704. doi:10.1038/s41467-019-12725-1

28. Akam T, Rodrigues-Vaz I, Marcelo I, et al. The Anterior Cingulate Cortex Predicts Future States to Mediate Model-Based Action Selection. Neuron. 01 06 2021;109(1):149–163.e7. doi:10.1016/j.neuron.2020.10.013

29. White JK, Bromberg-Martin ES, Heilbronner SR, et al. A neural network for information seeking. Nat Commun. Nov 14 2019;10(1):5168. doi:10.1038/s41467-019-13135-z

30. Rushworth MF, Behrens TE. Choice, uncertainty and value in prefrontal and cingulate cortex. Nat Neurosci. Apr 2008;11(4):389–97. doi:10.1038/nn2066

31. Behrens TE, Woolrich MW, Walton ME, Rushworth MF. Learning the value of information in an uncertain world. Nat Neurosci. Sep 2007;10(9):1214–21. doi:10.1038/nn1954

32. Hooks BM, Papale AE, Paletzki RF, et al. Topographic precision in sensory and motor corticostriatal projections varies across cell type and cortical area. Nat Commun. Sep 03 2018;9(1):3549. doi:10.1038/s41467-018-05780-7

33. Frank MJ. Hold your horses: a dynamic computational role for the subthalamic nucleus in decision making. Neural Netw. Oct 2006;19(8):1120–36. doi:10.1016/j.neunet.2006.03.006

34. Kaufman MT, Churchland MM, Ryu SI, Shenoy KV. Vacillation, indecision and hesitation in moment-by-moment decoding of monkey motor cortex. Elife. May 05 2015;4:e04677. doi:10.7554/eLife.04677

35. Peters AJ, Fabre JMJ, Steinmetz NA, Harris KD, Carandini M. Striatal activity topographically reflects cortical activity. Nature. Mar 2021;591(7850):420–425. doi:10.1038/s41586-020-03166-8

36. Kravitz AV, Owen SF, Kreitzer AC. Optogenetic identification of striatal projection neuron subtypes during in vivo recordings. Brain Res. May 20 2013;1511:21–32. doi:10.1016/j.brainres.2012.11.018

